## Supplementary Figures 1-8 and Tables 3-5 for "Allelic expression analysis of Imprinted and X-linked genes from bulk and single-cell transcriptomes"

#### Supplementary Fig. 1

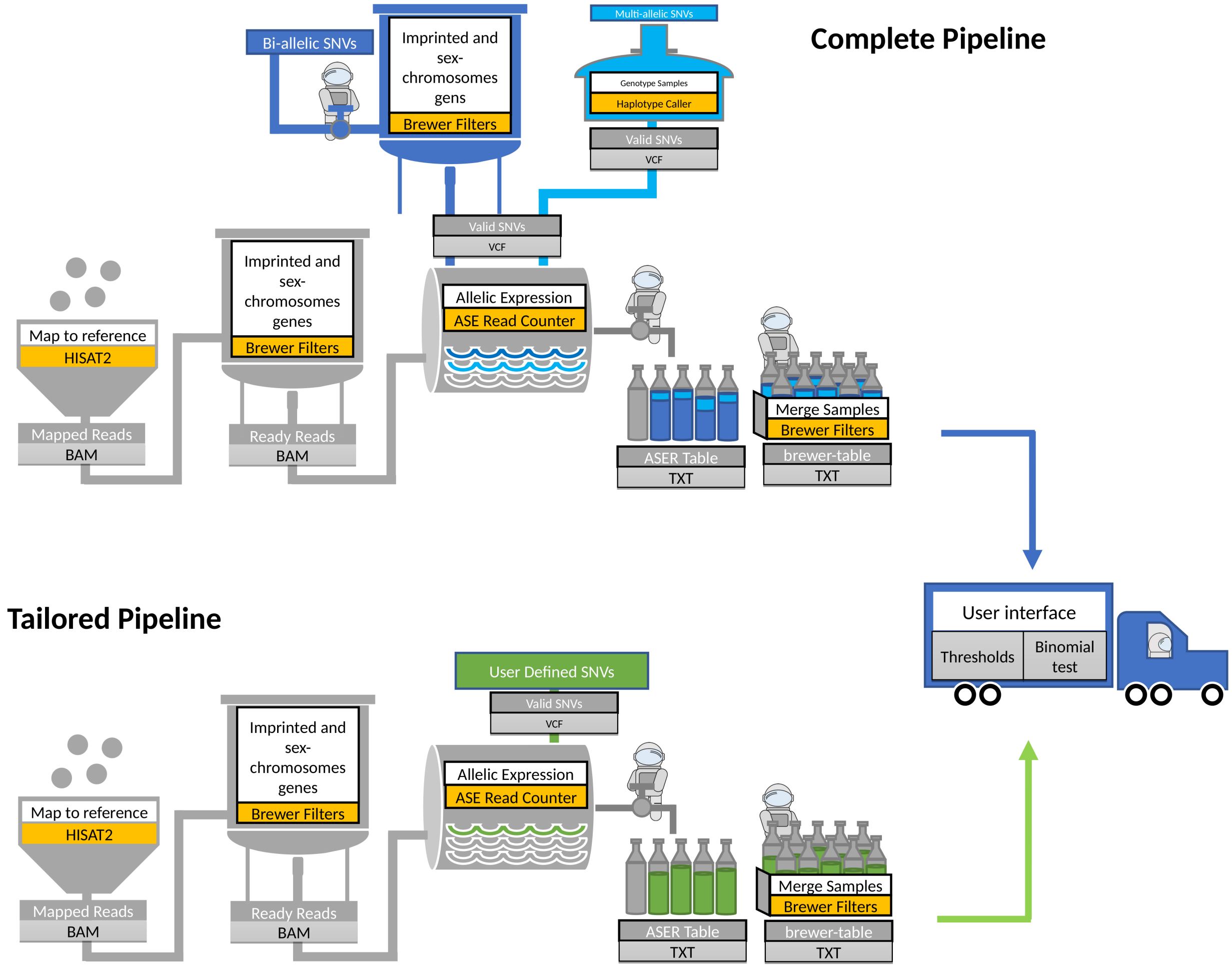


**Supplementary Fig. 1.** Complete and Tailored pipelines overview. The Complete pipeline sacrifices speed for the sake of completeness by using a larger set of SNVs obtained merging i) the SNVs called by HaplotypeCaller (https://gatk.broadinstitute.org) on the user dataset using a pre-compiled set of multi-allelic SNVs and ii) the bi-allelic set used in the Standard pipeline. The use of a larger set of SNVs will increase the power to detect bi-allelic expression. The Tailored Pipeline is meant for users that need to evaluate their own set of SNVs for example those computed using DNA-seq data of matched samples. This allows the user to evaluate imprinting and X-inactivation starting directly from the actual SNV profile of the samples. While the input files for the Standard and the Complete pipelines are only fastq files derived from RNA-seq experiments, the Tailored pipeline additionally requires the VCF file with a set of bi-allelic SNVs. Both Complete and Tailored pipeline results can be further analysed and visualized through the User Interface.

**Supplementary Fig. 2**

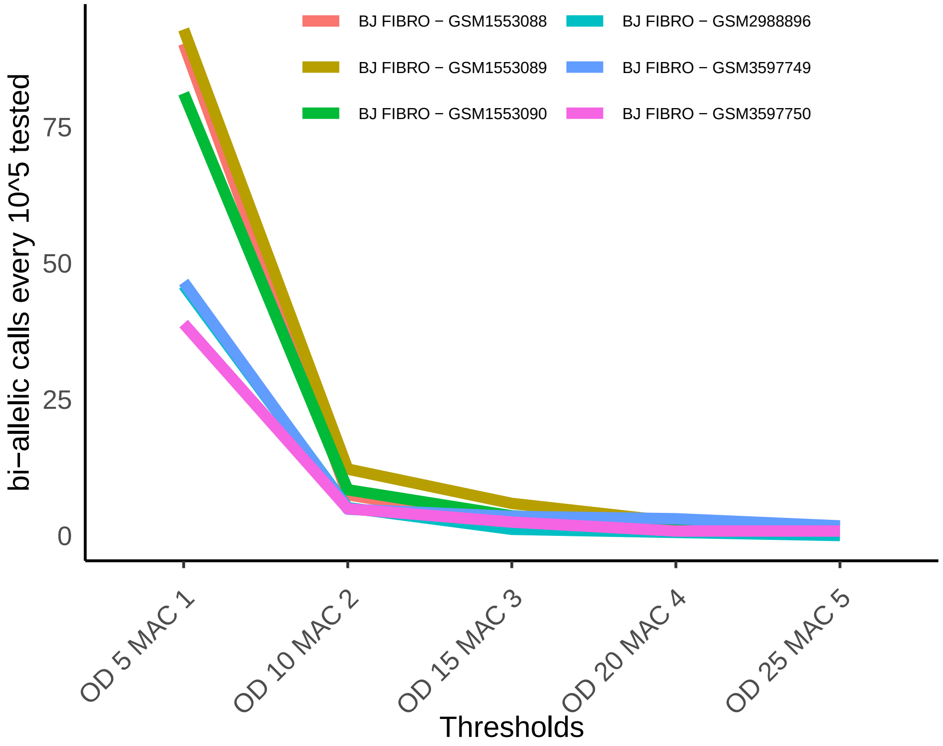


**Supplementary Fig. 2**. False positives bi-allelic calls estimated by analysis of transcripts on the X chromosome in 6 male BJ fibroblasts samples. On the x axis thresholds combination of overall depth (OD) and minor allele coverage (MAC).

**Supplementary Fig. 3**

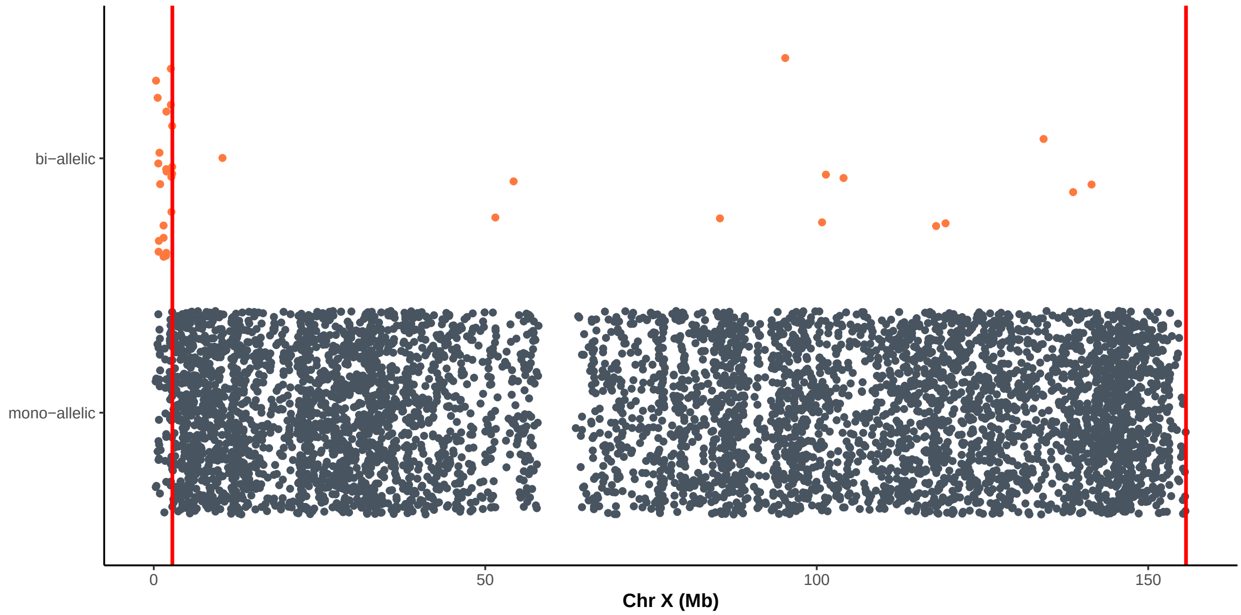


**Supplementary Fig. 3**. BJ fibroblasts genomic DNA analyzed by SNP Array (Affymetrix Mapping 250K Nsp SNP Array, GEO: GSM1868966, series GSE72531). In grey, all the mono-allelic SNVs. In red, bi-allelic SNVs. The vertical red lines indicate the boundaries of PseudoAutosomal Regions (PARs) at both tips of the X chromosome. Excluding the PARs, we observed 13 bi-allelic SNVs out of 5444 analysed (2 every 1,000).

Supplementary Fig. 4

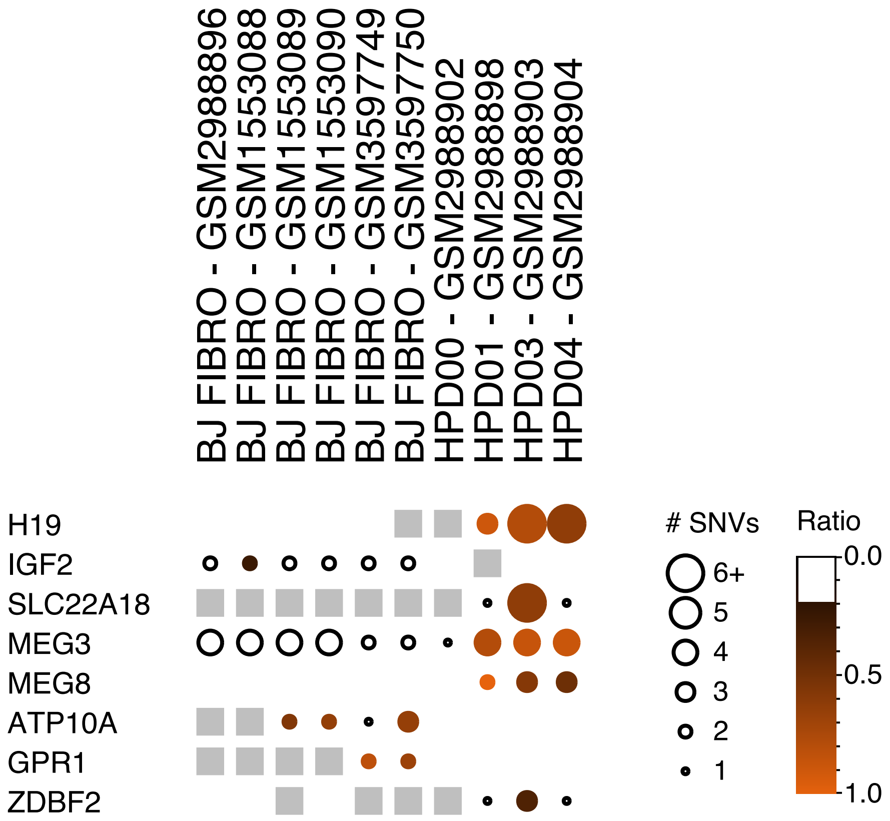


**Supplementary Fig. 4**. BrewerIX gene summary panel. Results on isogenic BJ fibroblasts and both primed and naive iPSCs obtained by BrewerIX using the Standard pipeline. The larger the dot, the higher the number of SNVs supporting the bi-allelic estimate, the darker the orange, the closer to 0.5 is the average of the ratios of all the bi-allelic SNVs. Empty spots are genes with no evidence of bi-allelic expression, white spots are genes with no read overlapping any SNVs

**Supplementary Fig. 5**

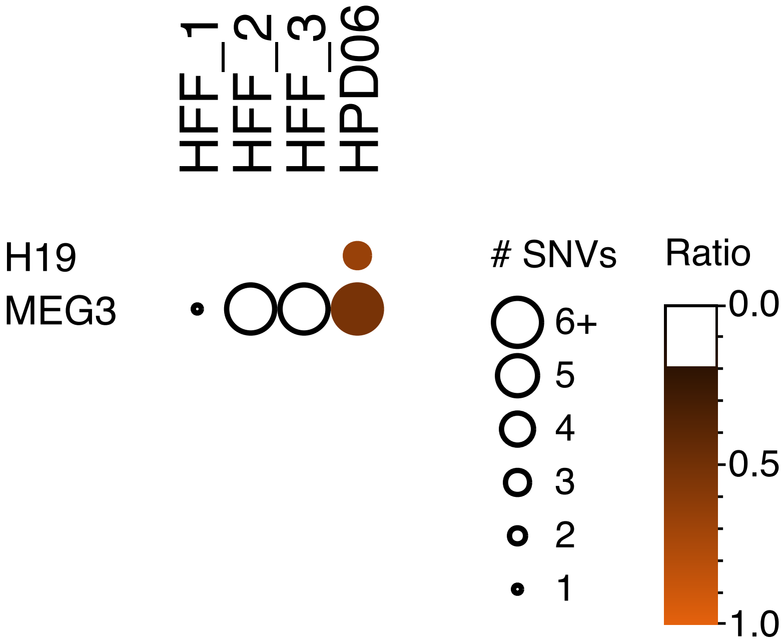


**Supplementary Fig. 5**. BrewerIX gene summary panel. Results on HFF iPS reprogramming dataset (GSE110377) and 3 normal HFF (GSE93226) obtained by BrewerIX in Standard mode. The larger the dot, the higher the number of SNVs supporting the bi-allelic estimate, the darker the orange, the closer to 0.5 is the average of the ratios of all the bi-allelic SNVs. Empty spots are genes with no evidence of bi-allelic expression, white spots are genes with no read overlapping any SNVs.

### Supplementary Fig. 6

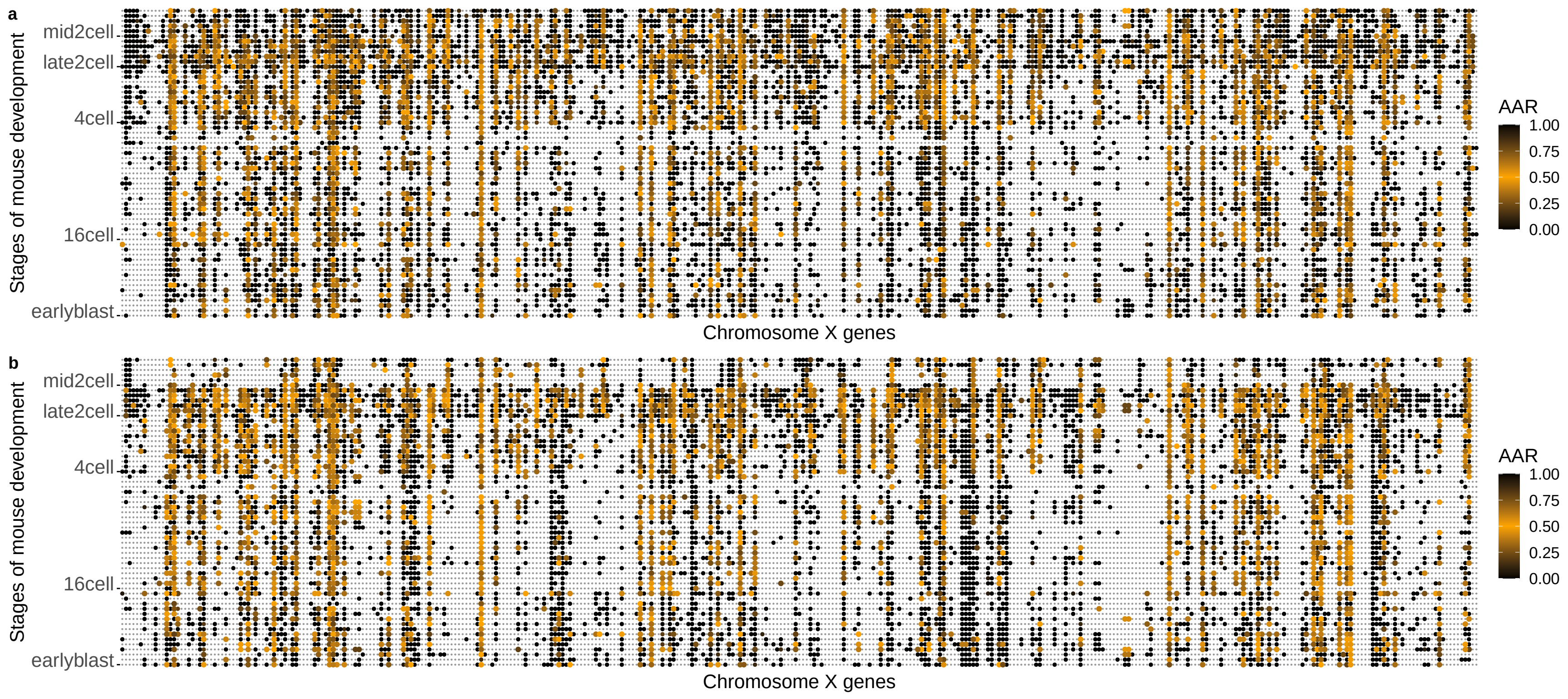


**Supplementary Fig. 6**. X-chromosomes activation status of female samples from dataset GSE45719. Single cells from early mouse embryos at the indicated stage were analysed by single-cell RNA sequencing. a, Average Allelic Ratio of X-chromosome genes (AAR, maternal / total) from the custom analysis performed by Deng and colleagues. b, Average Allelic Ratio of X-chromosome genes (AAR, minor / tot) using BrewerIX computed values. Black dots indicate mono-allelic expression; orange dots indicate bi-allelic expression (AAR=0.5). Grey dots indicate transcripts that were not detected.

#### Supplementary Fig. 7

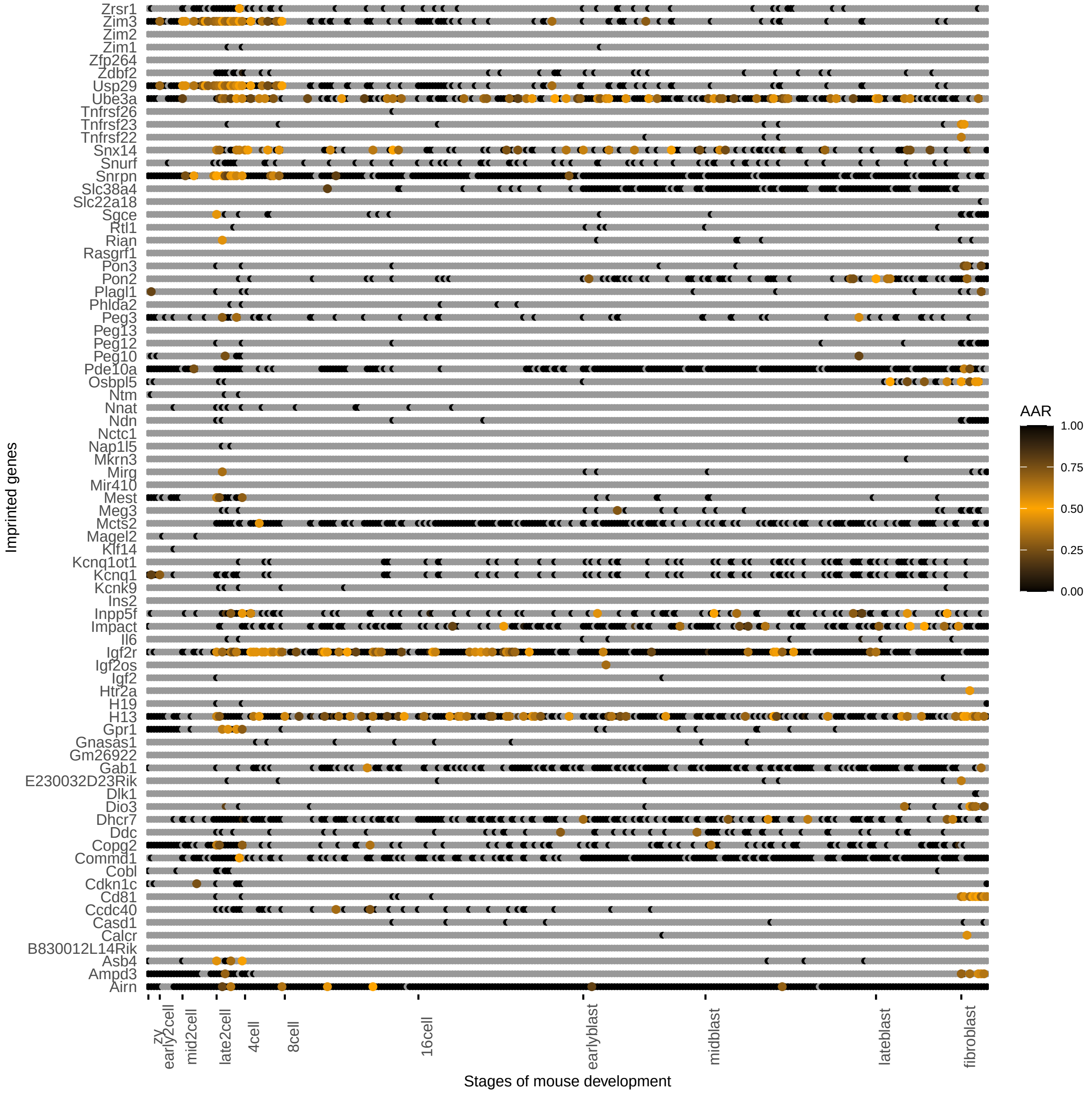


**Supplementary Fig. 7**. Average allelic ratio of imprinted genes across developmental stages computed by BrewerIX on dataset GSE45719. Black dots indicate mono-allelic expression; orange dots indicate bi-allelic expression (AAR=0.5). Grey dots indicate transcripts that were not detected. See also Figure 2c.

#### Supplementary Fig. 8

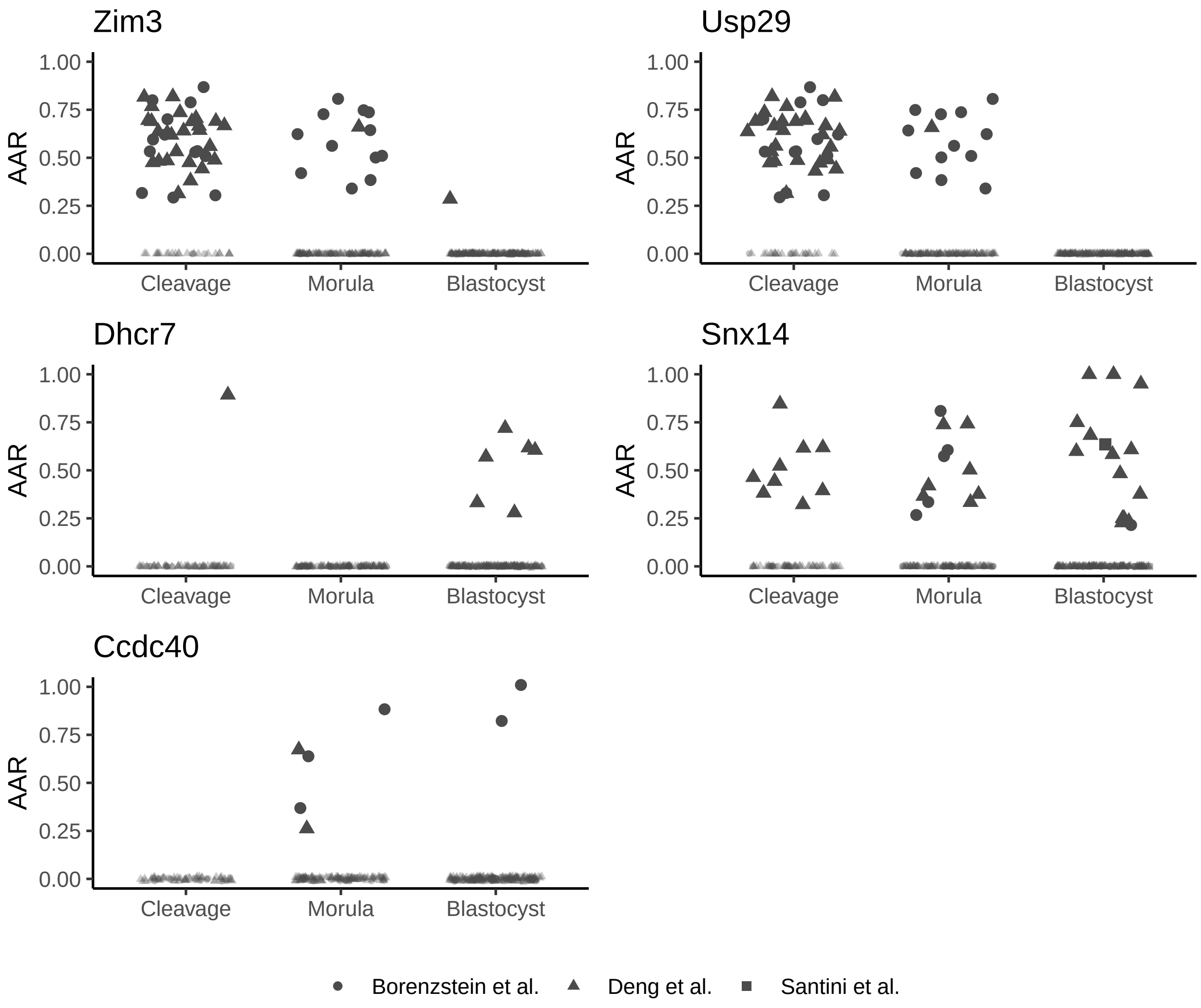


**Supplementary Fig. 8**. Genes with frequent LOI in mouse embrio development obtained by studying three dataset (Borensztein et al., Santini et al., and Deng et al.). On the y axis, the Average Allelic Ratios (AAR) of single samples (single cells or bulk for Santini et al dataset). Developmental stages has been collapsed into broader categories (Cleavage, Morula and Blastocyst).

#### Table S1

**Table S1**. Human manually curated imprinted genes. Three resources were merged: geneimprint (http://geneimprint.com/), Otago (http://igc.otago.ac.nz/home.html) and Santoni FA et al., Am J Hum Genet. 2017 (https://doi.org/10.1016/j.ajhg.2017.01.028). Isoform dependent imprinted genes are marked along with their source database. Placental imprinted genes are labeled following Okae et al. 2014 (https://doi.org/10.1371/journal.pgen.1004868)

#### Table S2

**Table S2**. Mouse manually curated imprinted genes. Three resources were merged: geneimprint (http://geneimprint.com/), Otago (http://igc.otago.ac.nz/home.html) and Inoue A et al., Nature 2017 (https://doi.org/10.1038/nature23262). Isoform dependent imprinted genes are marked along with their source database. Placental imprinted genes are labeled following Okae et al. 2012 (http://10.1093/hmg/ddr488)

#### Table S3

**Table S3**. List of datasets used for the case studies. Details on number of samples, organism, figure panels where the dataset results are shown, samples' types (bulks or single-cell RNAseq), average sequencing depth per samples, BrewerIX pipeline used and run times on a Linux Gnome Desktop (12 CPU, 32 Gb Ram PC) are reported.

| Description | Datasets | Shown | Species | Type | Samples (n) | Average Depth | Run time Standard mode | Run time Complete mode |
|---|---|---|---|---|---|---|---|---|
| Parameter settings - BJ Fibroblasts | GSE110377; GSE126397; GSE63577 | Fig. 1b, Supplementary Fig 2 | Human | Bulk | 6 | 37 M | 2.5h | 3.5h |
| Reprogramming of BJ fibroblasts | GSE110377; GSE126397; GSE63577 | Fig. 1c,d, Supplementary Fig. 4 | Human | Bulk | 10 | 28 M | 3h | 4.5h |
| Reprogramming of HFF Fibroblasts | GSE93226; GSE110377 | Supplementary Fig. 5 | Human | Bulk | 3 | 21 M | 1h | 1.5h |
| mESCS - 2i/L and S/L conditions | GSE84164 | Fig. 1f | Mouse | Bulk | 8 | 22M | 2h | 3h |
| mESCs - bulk and single-cell RNAsea | E-MTAB-2600 | Fig. 1g, Fig 2a | Mouse | SC / Bulk | 672/3 | 7 M [70M - 0.004M] / 52M | 16h | - |
| Early mouse development | GSE45719 | Fig. 2b,c, Supplementary Fig. 6-7-8 | Mouse | SC | 296 | 22M [139M - 1.3M] | 7h | - |
| Blastocyst-stage embryos | GSE152106 | Fig 2c, Supplementary Fig. 8 | Mouse | Bulk | 8 | 35M [43.9M - 25.3M] | | 1h |
| Early mouse development | GSE80810 | Fig 2c, Supplementary Fig. 8 | Mouse | SC | 113 | 18M [85M - 6M] | | 5h |
| Human fibroblasts and lymphoblastoid cells | GSE123028 | Fig. 2d,e | Human | SC | 820 | 18 M [37M - 0.0004M] | 20h | - |
| Breast cancer- bulk and single-cell RNAseq | GSE75688 | Fig. 2g | Human | SC / Bulk | 515/12 | 5.8M [11.1M - 3.1M] / 10M [7.8M - 4.6M] | 5h (SC only) | 2.5h (bulk only) |
| Fetal neocortex and human cerebral organoids | GSE75140 | Fig 3, Supplementary Fig 9 | Human | SC | 734 | 1.95M [4.02M - 0.01M] | 11h | - |
| ESC-derived MiniBrains | GSE124174 | Fig 3, Supplementary Fig 9 | Human | SC | 3 | 20.3M; 22.2M; 18.9M | - | 1h |
| iPSC-derived MiniBrains | GSE86153 | Fig 3, Supplementary Fig 9 | Human | SC | 2 | 301M; 250M | - | 3h |
| ESC and iPSC derived cortical organoids | E-MTAB-8337 | Fig 3, Supplementary Fig 9 | Human | SC | 2 | 66.9M; 50.6M | - | 2h |
| iPSC derived cortical organoids | GSE112137 | Fig 3, Supplementary Fig 9 | Human | Bulk | 4 | 23.5M; 10.1M; 9.5M; 26.0M | - | 3.5h |
| ESC and iPSC derived cortical organoids | E-MTAB-8325 | Fig 3, Supplementary Fig 9 | Human | Bulk | 4 | 25.9M; 17.7M; 35.0M; 29.0M | - | 2h |

#### Table S4

**Table S4**. List of SNVs validated by PCR followed by Sanger sequencing. Chromosome position, gene symbol, reference and alternative alleles along with their read counts for each sample are reported.

| Validation of SNVs by PCR followed by Sanger sequencing | | | | | | | | HPD01 Naive iPSCs - GSM2988898 | | HPD03 Naive iPSCs - GSM2988903 | | HPD04 Naive iPSCs - GSM2988904 | |
|---|---|---|---|---|---|---|---|---|---|---|---|---|---|
| **SNV** | **Chr** | **position** | **ref** | **alt** | **Gene symbol** | **Sanger sequencing** | **Expression in naive iPSCs (HPD03)** | **Ref** | **Alt** | **Ref** | **Alt** | **Ref** | **Alt** |
| rs2228613 | 19 | 10154917 | G | T | DNMT1 | confirmed | bi-allelic | 23 | 7 | 14 | 33 | 17 | 10 |
| rs61750052 | 19 | 10154368 | G | A | DNMT1 | confirmed | bi-allelic | 22 | 7 | 26 | 19 | 16 | 13 |
| rs2839703 | 11 | 1995432 | T | C | H19 | confirmed | bi-allelic | 101 | 91 | 92 | 19 | 136 | 57 |
| rs2839704 | 11 | 1995429 | T | C | H19 | confirmed | bi-allelic | 102 | 91 | 92 | 20 | 132 | 56 |
| rs3741219 | 11 | 1995389 | A | G | H19 | confirmed | bi-allelic | 51 | 47 | 51 | 51 | 79 | 52 |
| rs2400941 | 14 | 100834230 | C | G | MEG3 | confirmed | bi-allelic | 142 | 115 | 457 | 460 | 150 | 166 |
| rs8013873 | 14 | 100835753 | C | T | MEG3 | confirmed | bi-allelic | 382 | 267 | 1128 | 886 | 335 | 271 |
| rs2289998 | 11 | 3088178 | C | T | OSBPL5 | confirmed | bi-allelic | - | - | 3 | 2 | - | - |
| rs2289999 | 11 | 3088173 | A | G | OSBPL5 | confirmed | bi-allelic | - | - | 3 | 2 | - | - |
| rs935431 | 11 | 3088067 | G | A | OSBPL5 | confirmed | bi-allelic | - | - | 4 | 1 | - | - |
| rs11555134 | 7 | 50591496 | C | T | GRB10 | confirmed | biallelic | 2 | 5 | - | - | 3 | 2 |
| rs2192206 | 15 | 23686360 | G | A | NDN | confirmed | bi-allelic | 18 | 9 | 18 | 13 | 27 | 15 |

#### Table S5

**Table S5**. Primers used for PCR and Sanger sequencing

| Primer Name | Sequence (5' -> 3') |
|---|---|
| DNMT1__rs61750052f | CGGCCTCATCGAGAAGAATA |
| DNMT1__rs61750052r | GTGATCCTCTGGCCTCAGAC |
| DNMT1__rs2228613f | TCAGCAAGATTGTGGTGGAG |
| DNMT1__rs2228613r | CCAAACTGGCCTAAATCCAA |
| GRB10__f | TGGCCATTTCCTTACCTTTG |
| GRB10__r | GGGTTGACTGAGGAGCAGAG |
| H19__f | TTCAAAGCCTCCACGACTCT |
| H19__r | GTCGGAGCTTCCAGACTAGG |
| MEG3__rs2400941f | CGGGTGTAGACCTCTGAAGC |
| MEG3__rs2400941r | TTTGGGGCCTGTATGTGAAT |
| MEG3__rs8013873f | GCCCTCCTGTGGTCTGAGTA |
| MEG3__rs8013873r | ACGATCACGAGGGGTCTCT |
| NDN__f | CCCGAATACGAGTTCTTTTGG |
| NDN__r | TGAATACTGCACTGTAAATCCTGAA |
| OSBPL5__f | ACCACATCCTCAAATAGGAG |
| OSBPL5__r | CCTGCAGAGAGGCCAGTG |
